# Spatially-Resolved Proteomic Analysis of the Lens Extracellular Diffusion Barrier

**DOI:** 10.1101/2021.02.23.432581

**Authors:** Zhen Wang, Kevin L Schey

## Abstract

**Purpose:** The presence of a physical barrier to molecular diffusion through lenticular extracellular space has been repeatedly detected in multiple species. This extracellular diffusion barrier has been proposed to restrict the movement of solutes into the lens and to direct nutrients into the lens core via the sutures at both poles. The purpose of this study is to characterize the molecular components that could contribute to the formation of this barrier.

**Methods:** Three distinct regions in the bovine lens cortex were captured by laser capture microdissection guided by dye penetration. Proteins were digested by endoproteinase Lys C and trypsin. Mass spectrometry-based quantitative proteomic analysis followed by gene ontology (GO) and protein-protein interaction network analysis was performed.

**Results:** Dye penetration showed that lens fiber cells first shrink the extracellular spaces of the broad sides of fiber cells followed by closure of the extracellular space between narrow sides at normalized lens distance (r/a) of 0.9. Accompanying the closure of extracellular space of the broad sides, dramatic proteomic changes were detected including up-regulation of several cell junctional proteins. AQP0 and its interacting partners ERM proteins were among a few proteins that were upregulated accompanying the closure of extracellular space of the narrow sides suggesting a particularly important role for the major lens membrane protein AQP0 in controlling the narrowing of the extracellular spaces between lens fiber cells. The results also provided important information related to biological processes that occur during fiber cell differentiation such as organelle degradation, cytoskeletal remodeling and GSH synthesis.

**Conclusions:** The formation of lens extracellular diffusion barrier is accompanied by significant membrane and cytoskeletal protein remodeling.

## Introduction

To achieve its normal function as an optical element and to maintain its transparency, the lens has evolved a unique structure that lacks light-scattering elements including blood vessels and cellular organelles (1). In addition, the terminally differentiated fiber cells are closely packed with the extracellular space smaller than the wavelength of the light to reduce light scattering (1). In the absence of a blood supply, this large tissue cannot rely on simple diffusion to deliver nutrients to the center of the lens. It has been demonstrated that the lens establishes a microcirculation system to deliver nutrients, remove wastes, maintain ionic homeostasis, and to control the volume of the fiber cells (1-3, 4). Based on the microcirculation system model, a circulating current, carried primarily by Na^+^ ions, directs nutrients extracellularly to the center of the lens at both poles via the sutures. Na^+^ ions along with water and metabolites in the center of the lens diffuse toward the lens equator intercellularly, i.e. from cell to cell across cell membranes, and flow toward the lens surface through gap junctions. The formation of the microcirculation system relies on spatially distinct distributions of ion channels and transporters including highly concentrated activity of sodium-potassium pumps in the anterior epithelial cells (5–6) and a high gap junction coupling conductance at the equator (7). The circulating current at the surface of the lens has been confirmed experimentally (8). Magnetic resonance imaging (MRI) has allowed visualization of fluid fluxes of heavy water (D_2_O) that are directed into the lens at the poles and then move circumferentially toward the equator in the lens cortex (9). This study also identified a zone of restricted extracellular space diffusion (9). A physical barrier to extracellular space diffusion has been reported in lenses from different species (10–11) which is consistent with the observed reduction of extracellular space between fiber cells to avoid light scattering that has been confirmed by electron microscopy (12). This extracellular diffusion barrier has been proposed to restrict the movement of solutes into the lens and acts to direct nutrients into the lens core via the suture at both poles (9).

The molecular mechanism that controls closing of the extracellular space and formation of the extracellular diffusion barrier has not been studied. Tight junctions are normally believed to control paracellular permeability and provide a diffusion barrier (13). Occluded zones similar to tight junctions were described between lens fiber cells in an early EM study (12). Several well-known tight junction proteins can be detected in lens fiber cells such as ZO-1, junction adhesion molecule 3 (JAM3) and coxsackie and adenovirus receptor (Cxadr) (14, 15). However, due to lack of core proteins of tight junction occludins or claudins, tight junctions were reported to be absent in lens fiber cells (14, 15, 16). In the absence of claudins, lens fiber cells express LIM2; a member of the PMP-22/EMP/MP20/Claudin Family that shares a tetraspanin topology and the WGLWCC signature sequence in the first extracellular loop (17). The precise function of LIM2 is unknown, but an adhesive function (18) and a role in fiber cell junction formation and organization has been reported (19). Another unique structure present in lens fiber cells is the square array junction (20–22). AQP0 and its cleavage products were found to be enriched in square array junctions (20, 21). Previous electron microscopy data suggested that the reduction of lenticular intercellular space was correlated with the formation of square array and membrane undulation (20, 21). Thus, square arrays might potentially drive the formation of complicated membrane interdigitations and serve to maintain an extremely narrow extracellular space (20, 22). In addition, it has been reported that gap junction plaques on the broad side of the outer fiber cells could restrict penetration of larger solutes through extracellular space (9).

The purpose of this study was to investigate proteins that could regulate and control the closure of lens extracellular spaces. Previous studies mainly relied on immunohistochemistry, and confocal and electron microscopy (10, 11, 20, 22). Mass spectrometry-based quantitative proteomic analysis is a powerful discovery approach to examine global proteomic dynamics. We hypothesize that an in-depth spatially resolved proteomic study will reveal proteins and pathways that could be involved in controlling extracellular space and diffusion. In this paper, a dye penetration experiment defined three distinct regions within less than 1.5 mm distant from the lens capsule to inner cortex in the equatorial region of bovine lenses. These three regions were then isolated by laser capture microdissection and proteome changes across three regions were studied with high-throughput quantitative mass spectrometry. Our results showed that the extracellular diffusion barrier is formed in a stepwise manner accompanied with dramatic changes in fiber cell junctional proteins, membrane and cytoskeletal proteins.

## Materials and Methods

### Dye penetration and confocal imaging

Bovine lenses were extracted from fresh cow eyes (Light Hill Meats, Lynnville, TN) and incubated in M199 medium (Sigma, St. Louis, MO) that contained 1mg/ml Texas Red-dextran (10,000 MW, Lysine Fixable, Thermo Fisher Scientific, Rockford, IL) at 37°C for 6-24 hr. All lenses used in this study were from cows of about 2-years-old. After incubation, lenses were rinsed with M199 media and fixed in 50 mL of 2% paraformaldehyde containing 0.01% glutaraldehyde for 4 days. The lenses were washed three times with PBS for 20 min and then cryoprotected with 10% sucrose in PBS for one day at 4°C, 20% sucrose for 5 hr at room temperature and 30% sucrose for one hour at room temperature and additional two days at 4°C. For sectioning, lenses were soaked in Tissue-Tek O.C.T. compound for 20 min and mounted in equatorial orientation on chucks and frozen in liquid nitrogen. Lenses were sectioned to 30 μm thickness using a CM 3050 Cryostat (LEICA CM 3050S, Leica Microsystems Inc., Bannockburn, IL). The sections were transferred to glass slides and washed three times in PBS before mounting on glass slides with mounting media (Vector Laboratories, Burlingame, CA). Confocal imaging was performed using Zeiss LSM 710 Confocal Microscope (Carl Zeiss, Oberkochen, Germany) with a 20X objective. Airyscan imaging was performed with a confocal laser scanning microscope ZEISS LSM 880 (Carl Zeiss AG, Oberkochen, Germany) with a Plan-Apochromat 63X/1.46 Oil Corr M27 objective.

### Laser Capture Microdissection (LCM)

Bovine lenses were incubated with Texas Red-labeled dextran for 18 hr as described above. Lenses were rinsed with M199 media and frozen immediately at −80°C. 12 μm thickness fresh frozen sections were obtained using the same cryostat as mentioned above. Four lenses from four different animals were used to generate four biological replicates. The lenses were sectioned from anterior pole and 10 equatorial sections were collected from each lens at around 4.2 mm distance from the anterior pole. The sections were collected on PEN membrane slides (Carl Zeiss, Munich, Germany) and stored at −80°C. Before LCM, sections were dehydrated with 75% ethanol for 1 min and 100% ethanol for 1 min and air dried. The LCM procedure was conducted using a PALM UV Laser MicroBeam laser capture microdissection system (Carl-Zeiss, Oberkochen, Germany). Tissue regions were selected based on the measured distance from the lens capsule. Depending on the regions, tissue was cut at an energy level of 38-43 and catapulted into an 0.2 mL Eppendorf tube cap containing 25 μL HPLC-grade water. Normally 3-4×10^6^ µm^2^ tissue can be collected in one cap and 1.6-2.8×10^7^ µm^2^ tissue was collected from each region in total. Three different regions were captured based on confocal images from the dye penetration experiment. The first sample captured contains fiber cells 50-350 µm from the lens capsule followed by capture of a region 450-800 µm from the capsule for the second sample and finally a region 950-1250 µm from the capsule for the third sample. The three regions captured were named outer cortex 1 (OC1), outer cortex 2 (OC2) and inner cortex (IC). The IC region corresponds to the start of the barrier region. Five samples (collected in 0.2 mL Eppendorf tube caps) from each region were pooled into one tube and dried in a speedvac and deemed ready for further processing.

#### Sample preparation and LC-MS/MS analysis

20 µL of 100mM ammonium bicarbonate containing 2 M urea, 10 mM TCEP, pH 8.0 was added to each sample and the samples were incubated at 25°C for 45 min. 2 µL of 2-chloroacetamide (500 mM) was added to each sample to alkylate free cysteines. The samples were incubated at 25°C for 45 min. The samples were then centrifuged at 20,000g for 5 min and the supernatant was carefully separated from the pellets (not visible). The remaining pellets were washed with 20 µL 100 mM ammonium bicarbonate, 2 M urea, pH 8.0 for another three times and the supernatants were pooled as the soluble fraction (SF). The remaining pellets were called the insoluble fraction (ISF). 5 µL of each SF was diluted 7-fold in water and used for a Bradford protein assay (Thermo Fisher Scientific, Rockford, IL).

15 µg of total protein from each soluble fraction was used for enzyme digestion. 2 M urea in 100 mM ammonium bicarbonate was added to each sample to regulate the final volume to 100 µL. 100 µL of 100mM ammonium bicarbonate was then added to each sample to reduce the urea concentration to 1 M. The proteins were digested by endoproteinase Lys C (1:25 enzyme to protein ratio) for 2 hr at 37°C, followed by trypsin digestion (1:25 enzyme to protein ratio) at 37°C for 6 hr. For each insoluble fraction, 5 µL of acetonitrile (ACN), and 45 µL of 100 mM ammonium bicarbonate was added to each pellet. The samples were then digested by Lys C and trypsin as above. A protein assay was not done for ISF samples due to the low amount of protein present. The amount of enzyme used for digestion of the ISF was based on the total proteins measured in SF of the corresponding sample (1:200 enzyme to SF protein ratio). ISF samples were dried in a speedvac after digestion and reconstituted in 0.1% formic acid for StageTip cleaning. SF samples were acidified and cleaned by StageTips as described by Rappsilber et al. (23). The StageTips were made by placing a small portion of Empore solid phase extraction disks (3M) in a 200 µL pipette tip. An additional 1 mg of C18 (Phenomenex Jupiter resin, 5 µm mean particle size, 300 Å pore size) was loaded into the tip for SF samples and 0.5 mg of C18 was loaded into the tip for ISF samples. Peptides were eluted in 70% acetonitrile containing 0.1% formic acid. All samples after StageTip cleaning were dried in a speedvac and reconstituted in 0.1% formic acid. The final concentration of the SF is 0.1 µg/µL. The amount of proteins in the ISF was estimated to be around 1/15 of the SF based on base peak intensities in LC-MS/MS runs. So ISF samples were reconstituted in 0.1% formic acid and the volumes were 1/15 of the volume used for reconstituting corresponding soluble fractions. 500 ng of total protein for each sample was used for LC-MS/MS analysis.

For LC-MS/MS analysis, samples were separated on a one-dimensional fused silica capillary column (200 mm x 100 µm) packed with Phenomenex Jupiter resin (3 µm mean particle size, 300 Å pore size) coupled with an UltiMate 3000 RSLCnano system (Thermo Scientific, San Jose, CA). A 76-minute gradient was performed, consisting of the following: 2-72 min, 2-40% ACN (0.1% formic acid); 75-78 min, 45-90% ACN (0.1% formic acid) balanced with 0.1% formic acid. The flow rate was 350 nL/min. The eluate was directly infused into a Q Exactive Plus instrument (Thermo Scientific, San Jose, CA) equipped with a nanoelectrospray ionization source. The data-dependent instrument method consisted of MS1 acquisition (R=70,000) from m/z 350-1600, using an MS AGC target value of 3e6, maximum ion time of 60 ms followed by up to 20 MS/MS scans (R=17,500) of the most abundant ions detected in the preceding MS scan. The MS2 AGC target value was set to 5e4, with a maximum ion time of 100 ms, HCD collision energy was set to 27, dynamic exclusion was set to 10 s.

#### Data analysis

Raw LC-MS/MS data were loaded into MaxQuant software (http://maxquant.org/, version 1.6.6.0) (24) and searched against a UniProt bovine database (downloaded 12/19/2019) with contaminant proteins included using the Andromeda search algorithm within MaxQuant (25). The search was performed with trypsin specificity with a maximum of 2 missed cleavage sites. Methionine oxidation, asparagine deamidation and protein N-terminal acetylation were variable modifications (up to 2 modifications allowed per peptide); cysteine was assigned a fixed carbamidomethyl modification. Precursor mass tolerance was set at 5 ppm and the minimum peptide length was 6 amino acids, and maximum peptide mass was 4600 Da. A false discovery rate (FDR) of 1% was applied for both peptide and protein filtering and at least 2 unique razor peptides (the peptides assigned to the protein group with the largest number of total peptides identified) were used for protein identification. Quantification settings were as follows: re-quantify with a second peak finding attempt after protein identification has completed; match full MS1 peaks between runs within a 0.7 min retention time window. The label free quantitation (LFQ) algorithm in MaxQuant (24) was used for protein quantitation. Only razor and unique peptides were used for protein level quantitation. LFQ was performed between samples OC1 and OC2 as well as between OC2 and IC. Protein quantification in ISF and SF was done separately.

The LFQ intensities of proteins from the MaxQuant searches were imported into Perseus (V1.5.2.3) (26). Contamination and reverse identifications were filtered, and the remaining protein LFQ intensities were Log2 transformed. For quantification, only proteins that were present in at least 75% of replicates in at least one region were used and missing values were imputed with random numbers from a normal distribution (1.8, 0.3). Welch’s test in Perseus was used for identification of significantly changed proteins between regions (p<0.05). Pearson correlations were calculated though the multiscatter plot function in Perseus. Hierarchical clustering was completed based on Euclidean distance. Volcano plots were done in Perseus with more stringent parameters (FDR 0.01 and S0=0.5) for visualizing more significantly changed proteins between regions. Enriched GO terms and pathways were determined using Fisher’s exact test with Benjamini-Hochberg FDR 0.05 and total quantified proteins as the background. Statistical comparison of protein intensities for some junctional proteins across three different regions in ISF were also performed in GraphPad Prism 8. Protein-protein network analyses were generated by STRING 11.0 (27) using medium or high confidence.

## Results

### Characterization of the extracellular diffusion barrier in bovine lenses

Dye penetration images (Figure 1) show that Texas Red-labeled dextran (∼10kDa) penetrated the lens through the extracellular space and two distinct regions in the outer cortex were observed. From the capsule to 350-400 µm, the dye easily penetrated the lens and dye signal in this region did not change with longer incubation times. Fiber cells in this region have regular hexagonal shapes. Further into the lens, fiber cells became more compact and dye penetration between the broad sides of the fiber cells was inhibited and penetration through the narrow sides remained. Dye penetration through all extracellular spaces completely stopped at 850-900 µm from the capsule. Using three different lenses, the average normalized lens distances r/a (where (a) represents the lens radius and (r) represents the distance from the lens core to the region of interest) for dye penetration were 0.919±0.007 for 6-hour incubation, 0.913±0.004 for 9-hour incubation, 0.896±0.014 for 18-hour incubation, and 0.897±0.006 for 24-hour incubation. The normalized distances for 18 hr and 24 hr incubated lenses were not statistically different, indicating a barrier to extracellular diffusion starts at average normalized lens distance of 0.9 in the inner cortex of the lens.

**Figure 1:**
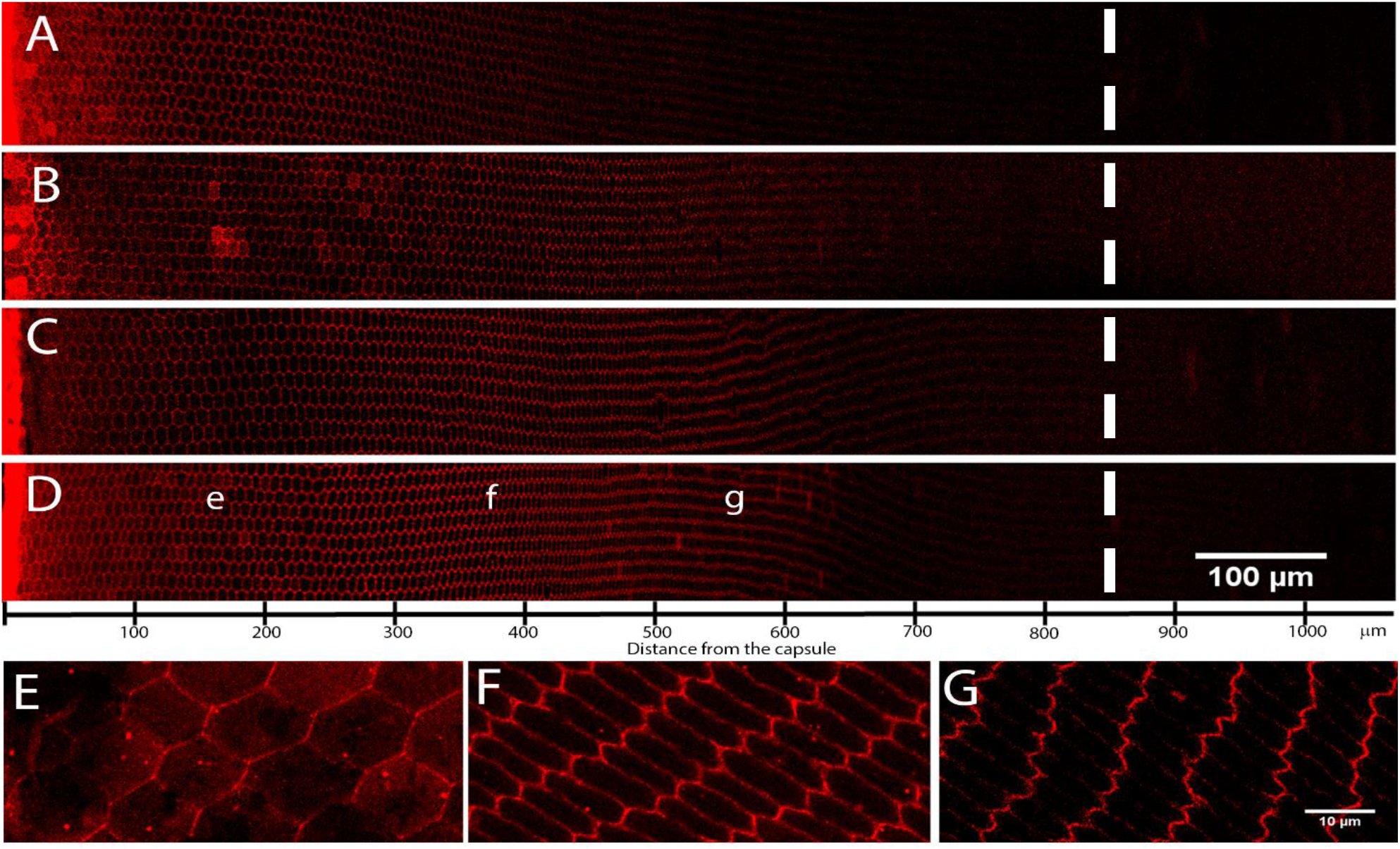
localization of the extracellular diffusion barrier. Bovine lenses were dissected from fresh cow eyeballs and incubated with lysine fixable Texas Red-dextran for various periods of time (6-24 hr). The lenses were then fixed and imaged by confocal microscopy (A-D). Dye incubation time was 6 hr (A), 9 hr (B), 18 hr (C) and 24 hr (D). The result showed the dye penetrated the lens through extracellular spaces and with further into the lens, dye penetration through the short sides of the fiber cells was inhibited. Regardless of incubation time, at 850-900 µm, dye penetration through extracellular spaces stopped. Airy scan images for corresponding regions in D were shown in E-G (corresponding to area labeled by e-g respectively) showing the fiber cell morphology and dye penetration changes in outer cortex of the lens. The dotted line indicates the position with normalized lens distance of 0.9.

### LFQ quantification shows dramatic proteome changes in lens cortex

Based on dye penetration experiments, three different regions in lens cortex were isolated by LCM. Note that the first 50 µm from the capsule was not captured to avoid the lens epithelium. The OC1 region (50-350 µm) represents peripheral fiber cells. In this region, the Texas Red-labeled dextran can penetrate through extracellular spaces surrounding the entirety of the fiber cells. The OC2 region (450-800 µm) represents the region where dye penetration through the broad sides of the fiber cells was inhibited. The IC region (950-1250 µm) represents the region where dye penetration is completely stopped. After LCM, the samples were separated into soluble and insoluble fractions and analyzed separately to comprehensively characterize soluble proteins and membrane and membrane associated protein expression by label free quantification (LFQ) using MaxQuant.

Data from three regions were initially searched together through MaxQuant and proteomes in different regions were found to be significantly altered. Normalization for quantitative proteomic studies relies on a majority of the proteome not being altered. The normalization step performed by MaxQuant has been shown to provide reliable protein quantification when one-third of the proteome was changing (24); however, to avoid potential effects of dramatic proteome changes on normalization, comparisons were only done between OC1 and OC2 regions or between OC2 and IC regions. The OC1 region was not compared to the IC region. In total, 1554 proteins were identified in the SF and ISF samples of OC1 and OC2. 1186 proteins were identified in SF and ISF of OC2 and IC. Venn diagrams showing the overlaps in protein identification among different samples are shown in Figure 2A and 2B. Even though Venn diagrams show significant overlaps in protein identification in different regions, a trend of a decrease in the number of proteins identified can be seen comparing OC2 to OC1 or IC to OC2. When protein signal intensities were compared, dramatic fiber cell proteome changes can be seen across these three regions that were derived from a narrow zone of the lens cortex. These dramatic proteome changes can be demonstrated by scatter plots (Figure 2 C and 2D) and Pearson correlations of protein signal intensities (Supplemental Figure 1). Pearson correlations between samples from different regions were consistently lower than correlations between biological replicates. Comparing the proteomes of SF and ISF from OC2 to OC1, scatter plots for average protein intensities of 4 biological replicates in different regions of the lens showed regional proteome changes in both SF and ISF (Figure 2C). Comparing the OC2 to the IC region, the proteome in SF showed less changes as shown in Figure 2D, but the ISF proteome showed more changes indicating dramatic changes of membrane and membrane associated proteins from OC2 to IC region.

**Figure 2:**
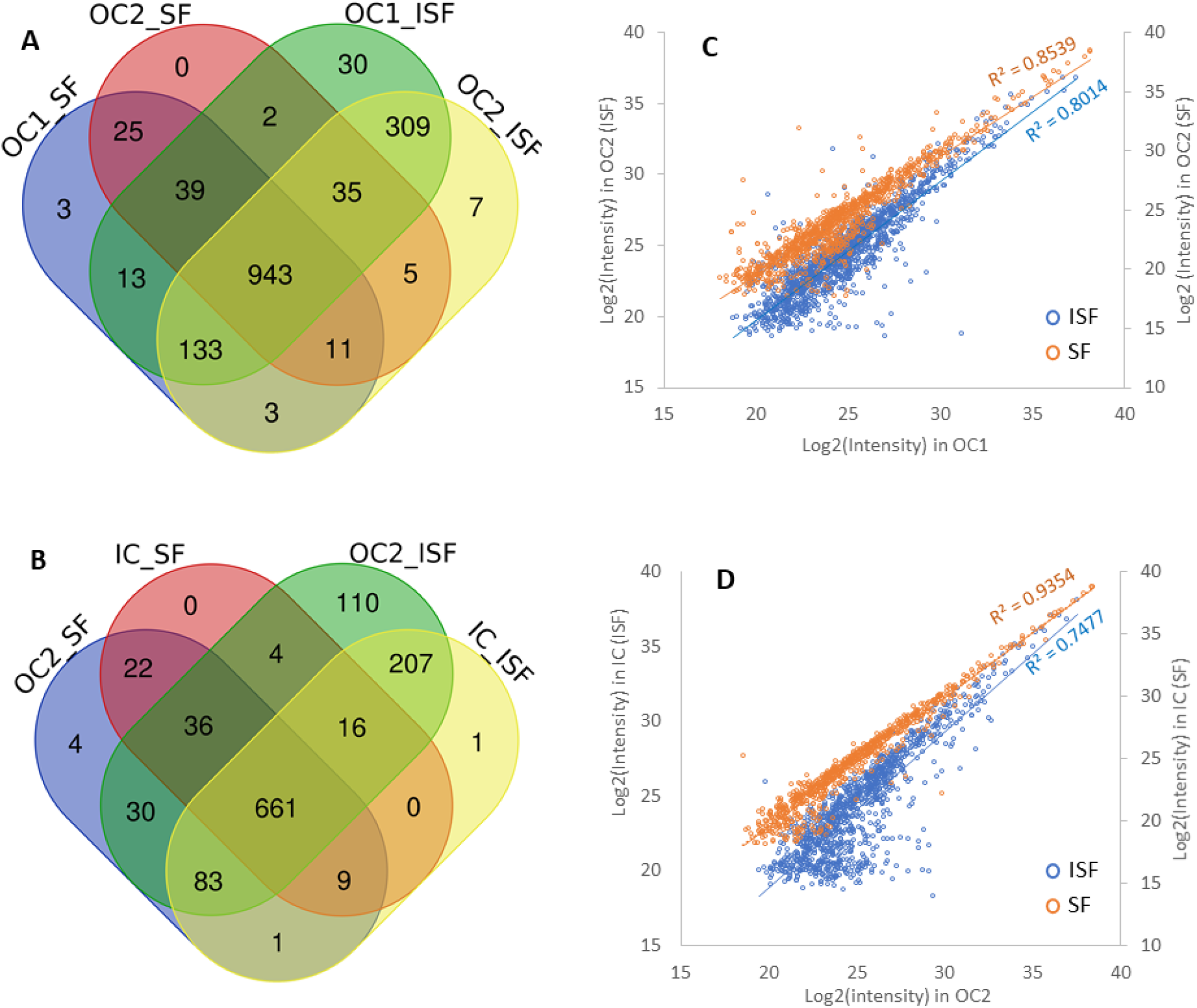
Venn diagrams and scatter plots. Three different regions of bovine lenses were isolated by LCM. The overlaps of the lists of proteins IDs detected by LC-MS/MS in samples of soluble fraction (SF) and insoluble fraction (ISF) from different regions (OC1, OC2, IC) of the lens are shown in Venn diagrams (A, B). Scatter plots of average protein intensities from 4 biological replicates are shown in C (OC1 vs OC2) and D (OC2 vs IC).

To study protein expression changes in the different regions of the lens, four Welch’s tests were performed (SF in OC1 vs SF in OC2; ISF in OC1 vs ISF in OC2; SF in OC2 vs SF in IC; ISF in OC2 vs ISF in IC). The results from four Welch’s tests were combined and are shown in Supplemental Table 1. A comparison of OC1 and OC2 results identified 250 proteins in SF and 306 proteins in ISF that were downregulated in OC2. 86 proteins in SF and 171 proteins in ISF were upregulated in OC2 compared with OC1. When comparing IC with OC2, 103 proteins in SF and 353 proteins in ISF were downregulated in IC and 71 proteins in SF and 133 proteins in ISF were upregulated in IC. Hierarchical clustering was then performed for statistically significantly changed proteins between regions and, in total, 8 clusters of proteins were identified. The results can be visualized by heat maps generated during clustering (Figure 3) and detailed information is included in Supplemental Table 1. Figures 3A and 3B show the heat maps generated from comparisons of ISF from different regions. As expected, the resulting dendrogram segregated different regions and 4 biological replicates clustered together. Therefore, these differentially expressed proteins provide molecular signatures that distinguish different regions of the lens cortex. Similar heat maps for proteins differentially expressed between regions in SF can be found in Supplemental Figure 2.

**Figure 3:**
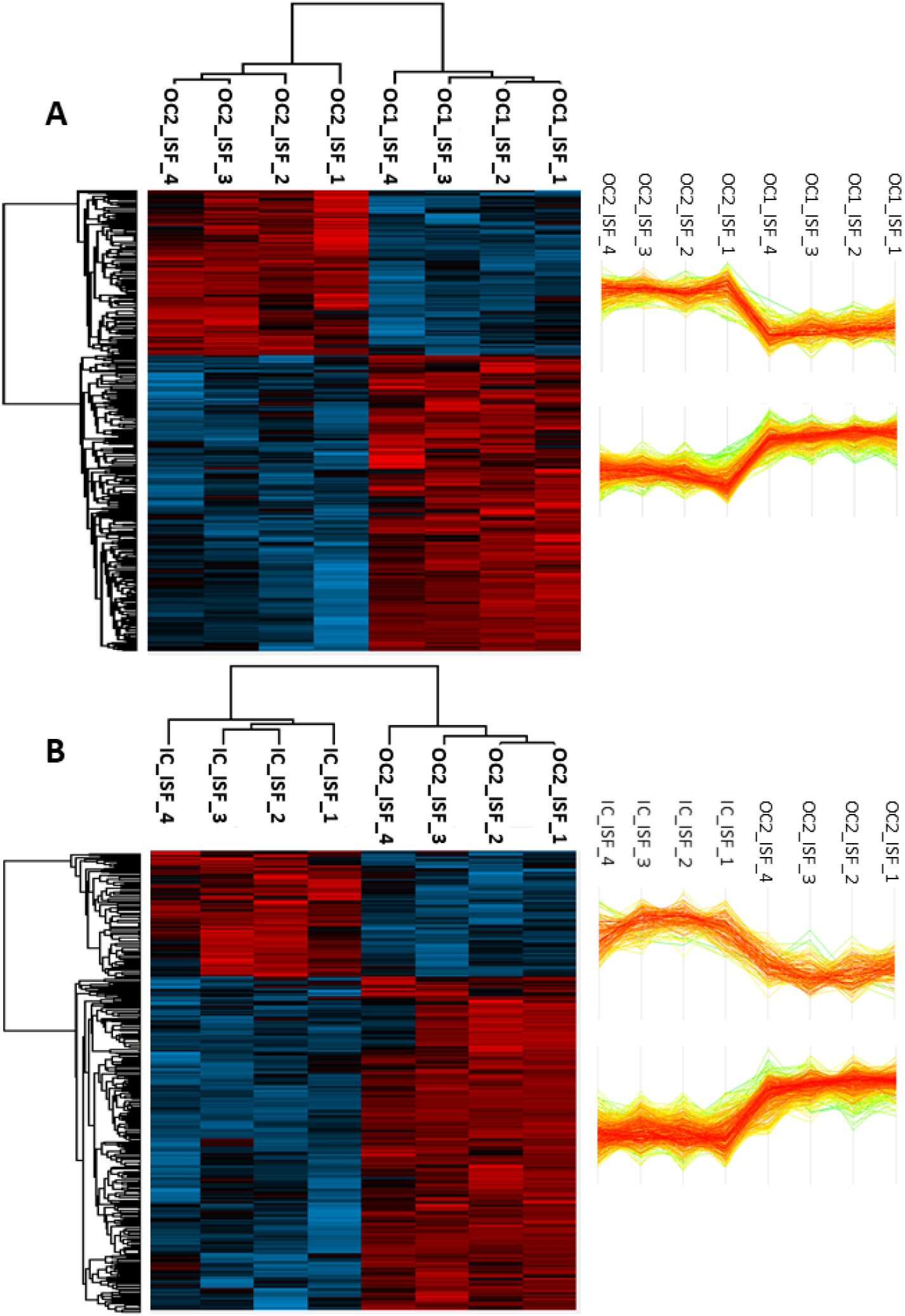
Heat map and hierarchical clustering of identified statistically significant changed proteins between different regions in ISF. Welch’s t-test was performed and hierarchical clustering was performed by Perseus based on Euclidean distance of statistically significant changed proteins (p<0.05). Hierarchical clustering defined two clusters of proteins for each comparison (A: comparing OC1 and OC2; B: comparing OC2 and IC). Profile plots shown on the right illustrate each protein intensity changes across all samples for a specific cluster.

To visualize proteins that show large magnitude changes with high statistical significance, volcano plots for ISF samples are shown in Figure 4. Corresponding volcano plots for SF samples can be found in Supplemental Figure 3. From OC1 to OC2, proteins showing a large fold increase or decrease can be detected (Figure 4A), whereas from OC2 to IC, many proteins significantly decrease in abundance and very few proteins dramatically increase abundance suggesting protein degradation in the IC region (Figure 4B). Proteins that show large fold changes include: cell-cell junctional proteins, cytoskeletal proteins, membrane proteins and some organelle-related proteins. The volcano plots show three proteins that are dramatically more abundant in the OC2 region compared to the OC1 region. These three proteins include lengsin (LGSN), chloride intracellular channel protein 5 (CLIC5), and lactase like protein (LCTL). Increasing expression of these proteins was detected in both SF (Supplemental Figure 3A) and ISF (Figure 4A). The abundance of these proteins is maintained in the IC region and does not show a significant change in the IC region compared to the OC2 region. Increased levels of CLIC5 are accompanied by a significant decrease of another chloride intracellular channel protein CLIC1 indicating a transition to a different chloride channel protein in fiber cells. In addition, several proteins (MAP7, EIF4A3 and SUMO2) showed large fold changes in only SF or ISF suggesting a change of solubility rather than a change in protein expression.

**Figure 4:**
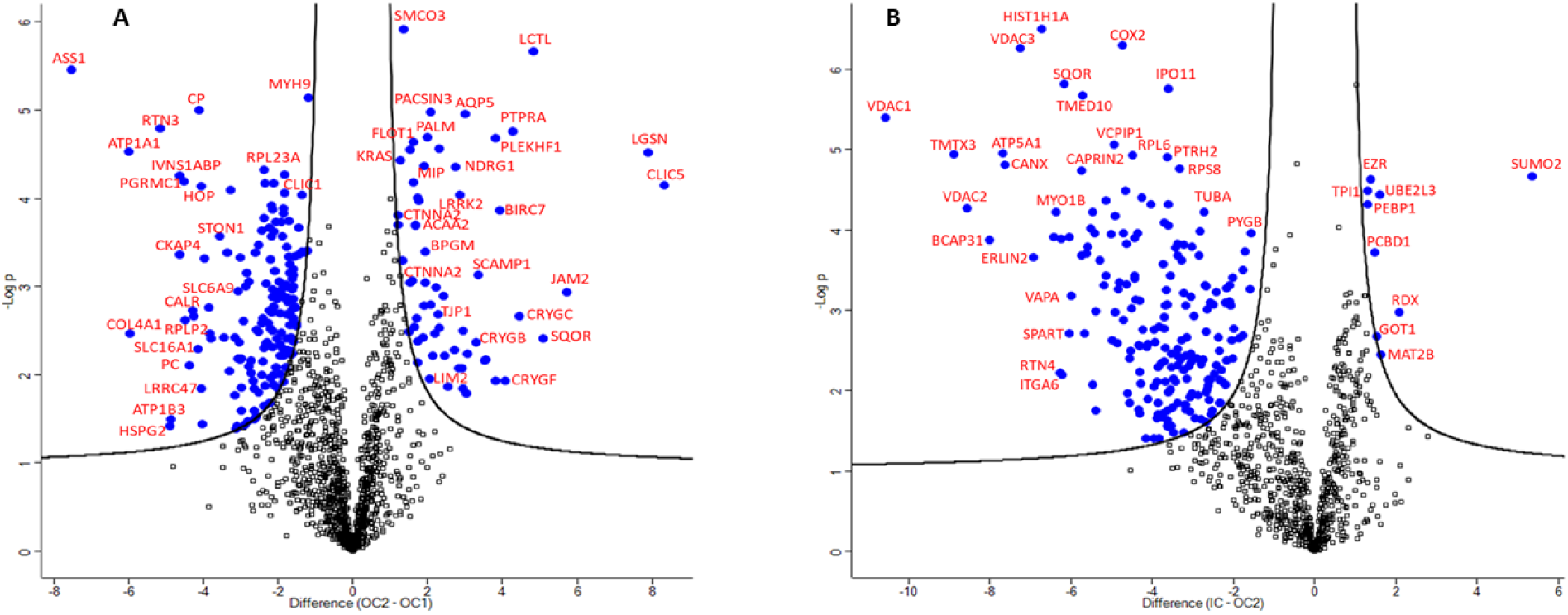
Volcano plots of spatially differentially expressed proteins measured in ISF. Proteins were graphed by fold change (difference) in x-axis and significance (-Log p) in y-axis using a false discovery rate of 0.01 and an S0 of 0.5. Statistical analysis was performed in Perseus. Blue circles represent proteins that are significantly changed between OC1 and OC2 (A) or OC2 and IC (B).

Gene Ontology (GO) enrichment analysis of differentially represented proteins against the background of all quantified proteins identified GO terms and pathways that are over- or under-represented in each region (Figure 5 and 6). As expected, in OC2, lens fiber cells have started to remove cellular organelles and reduce protein synthesis. GO terms such as ribosome, translation, endoplasmic reticulum lumen, had lower representation in the OC2 region compared to the OC1 region. Many proteins involved in these processes were not detected in IC samples or showed a downward trend in IC samples. Our data suggest mitochondrial proteins are degraded at a later stage of cell differentiation than ER and ribosomes. Most mitochondrial proteins did not decrease until cells were in the IC region. Many mitochondrial proteins increased in OC2 samples compared to OC1 samples, including proteins related to beta-oxidation (HADHA, HADHB, CPT1B, ETFDH) and some mitochondrial channel proteins (VDAC1, VDAC2, VDAC3, SLC25A13). Other than these expected protein changes related to organelle degradation, another over-represented pathway in OC2 samples is related to the proteasome and ubiquitination. This process was found to be further increased in IC samples. Glycolysis is over-represented in IC samples due to up-regulation of many enzymes involved in the glycolysis pathway (Supplemental Table 1). Another notable feature related to fiber cells in the IC region is the decrease of structural constituent cytoskeleton and microtubule including: filamins, spectrins, vimentin, ankyrins and tubulins, etc. Down-regulation of many cytoskeletal proteins was accompanied with up-regulation of several actin binding proteins such as Actin-related protein 2/3 complex subunit 1A and 3 (ARPC1A and ARPC3), F-actin-capping protein subunit alpha and beta (CAPZA2 and CAPZB), ezrin and radixin. Interestingly, glutathione synthase was found to be consistently increased in both SF and ISF in OC2 samples compared with OC1 samples and continued to increase in samples from the IC region. Another essential enzyme for GSH synthesis, glutamate-cysteine ligase catalytic subunit (GCLC), did not significantly change between regions. The biological process “response to reactive oxygen species” is significantly increased in IC samples in both SF and ISF including increased abundance of families of peroxiredoxin and glutathione S-transferase that could consume the reduced form of GSH. This result suggests that fiber cells may adapt a high GSH synthesis ability in the barrier region to fulfill the large GSH requirement of the lens.

**Figure 5:**
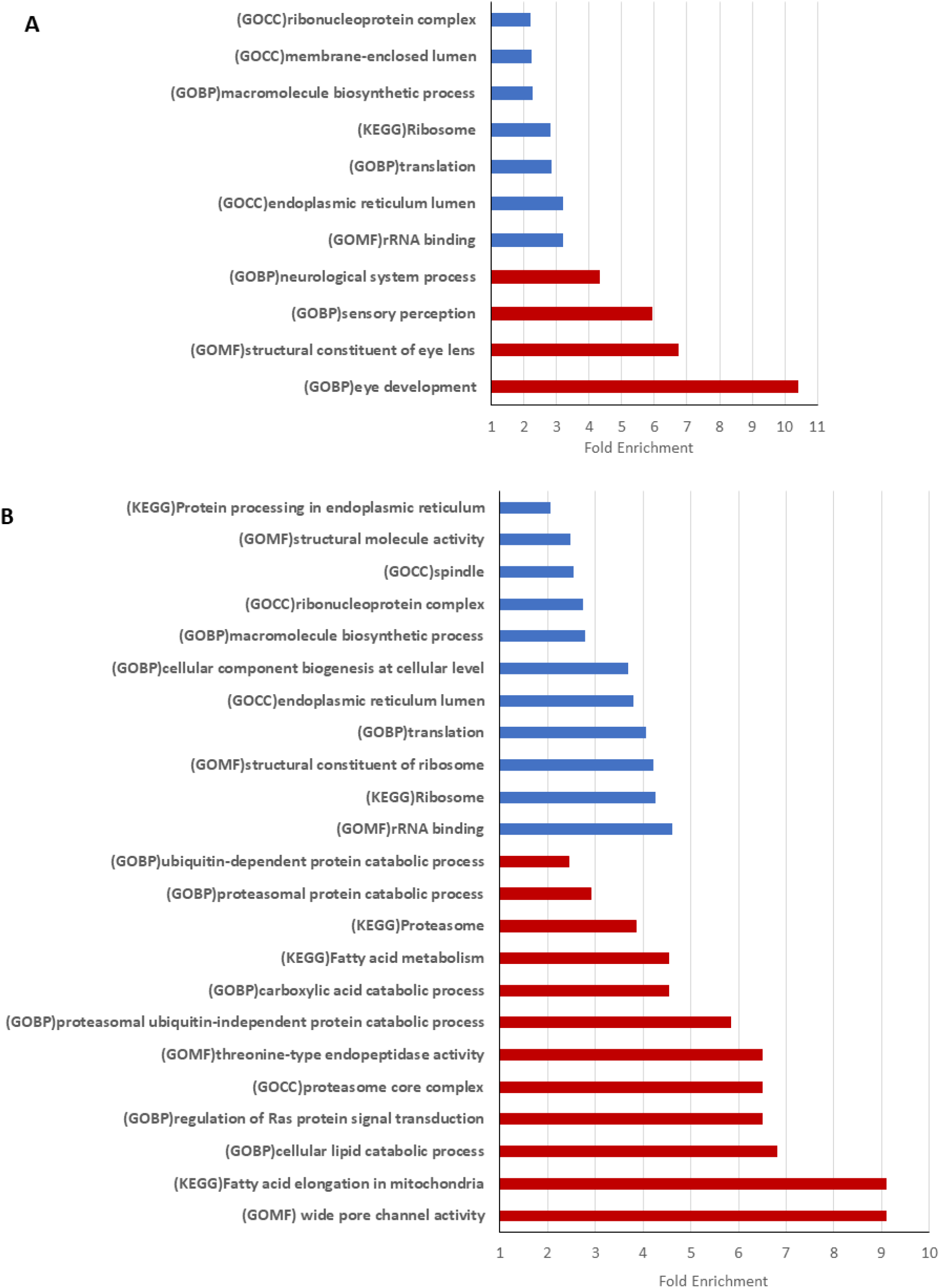
Go terms and pathway enrichment analysis for differentially expressed proteins in OC2 compared with OC1. Enriched GO terms and pathways were analyzed using a Fisher exact test with Benjamini-Hochberg FDR 0.05 and total quantified proteins as background. Only GO terms with fold change greater than 2 is shown. Blue and red bars represent GO terms and pathways that are decreased or increased respectively in OC2 compared with OC1. A shows enriched terms in SF and B shows enriched terms in ISF.

**Figure 6:**
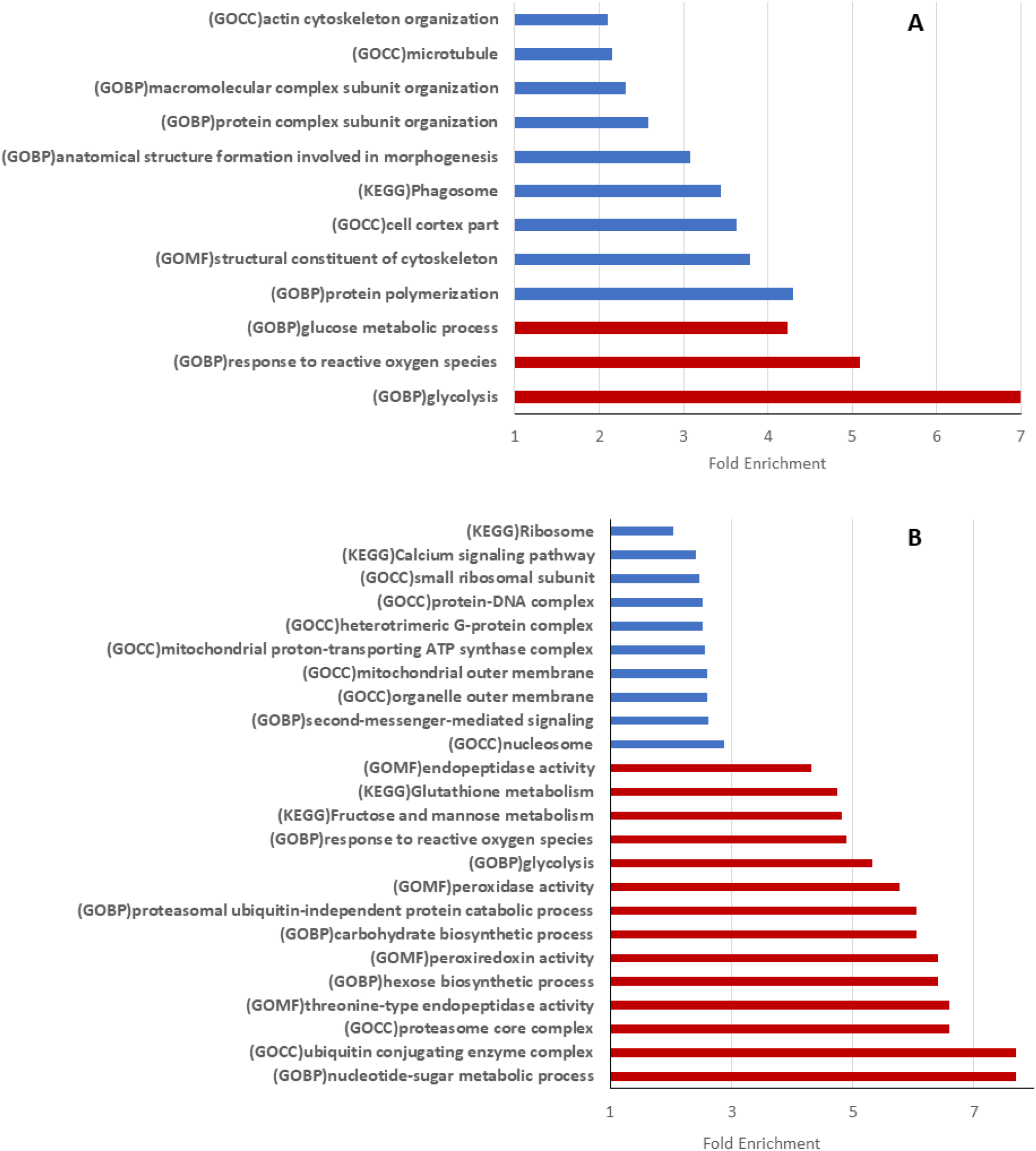
Go terms and pathway enrichment analysis for differentially expressed proteins in IC compared with OC2. Enriched GO terms and pathways were analyzed using a Fisher exact test with Benjamini-Hochberg FDR 0.05 and total quantified proteins as background. Only GO terms with fold change greater than 2 is shown. Blue and red bars represent GO terms and pathways that are decreased or increased respectively in IC compared with OC2. A shows enriched terms in SF and B shows enriched terms in ISF.

### The closing of extracellular diffusion is accompanied by changing of junction protein expression

A major goal of this study is to understand the molecular signatures that contribute to the closing of the extracellular space. Cell-cell junctions affect extracellular space and participate in paracellular barrier formation (28), therefore, proteins that are involved in cell-cell junctions were analyzed in captured lens regions. The results show that junction proteins are among the proteins that undergo significant changes between regions including proteins involved in adherens junctions, tight junctions, gap junctions, and square array junctions.

Adherens junctions and associated actin microfilaments are believed to be involved in stabilizing the structural integrity of lens cells during accommodation and in preserving a specific lens shape (29). Proteins involved in fiber cell adherens junctions reached maximum abundance in the OC2 region. These proteins include N-cadherin (CDH2), two isoforms of catenin alpha-2 (CTNNA2), catenin beta-1 (CTNNB1) and Armadillo repeat protein deleted in velo-cardio-facial syndrome (ARVCF). Two isoforms of CTNNA2 are among the most significantly changed proteins from OC1 to OC2 as shown in the volcano plot in Figure 4A. The increase of CTNNA2 coincided with a significant decrease of CTNNA1 (Supplemental Table 1), a result consistent with the transition from αE-catenin (epithelial type) to αN-catenin (neuronal type) with fiber cell differentiation reported previously (30). The decreasing of CTNNA1 was statistically significant in both ISF and SF samples, but more dramatic in SF samples (Supplemental Figure 3). To compare junction protein expression across three regions, LFQ intensities for these proteins across three different regions in the ISF are shown in Figure 7A. These data were obtained by MaxQuant searching of ISF samples from three regions together, therefore, the normalization was done across three regions. Again, statistical comparisons were only done between OC1 and OC2 regions and OC2 and IC regions. The results were similar to the previous analysis performed in Perseus. With the exception of CTNNA1, Figure 7A showed core proteins in adherens junction were higher in OC2 samples compared with OC1 or IC samples. The decrease in signal for lens adherens junction proteins in IC samples compared with OC2 samples was statistically significant.

**Figure 7:**
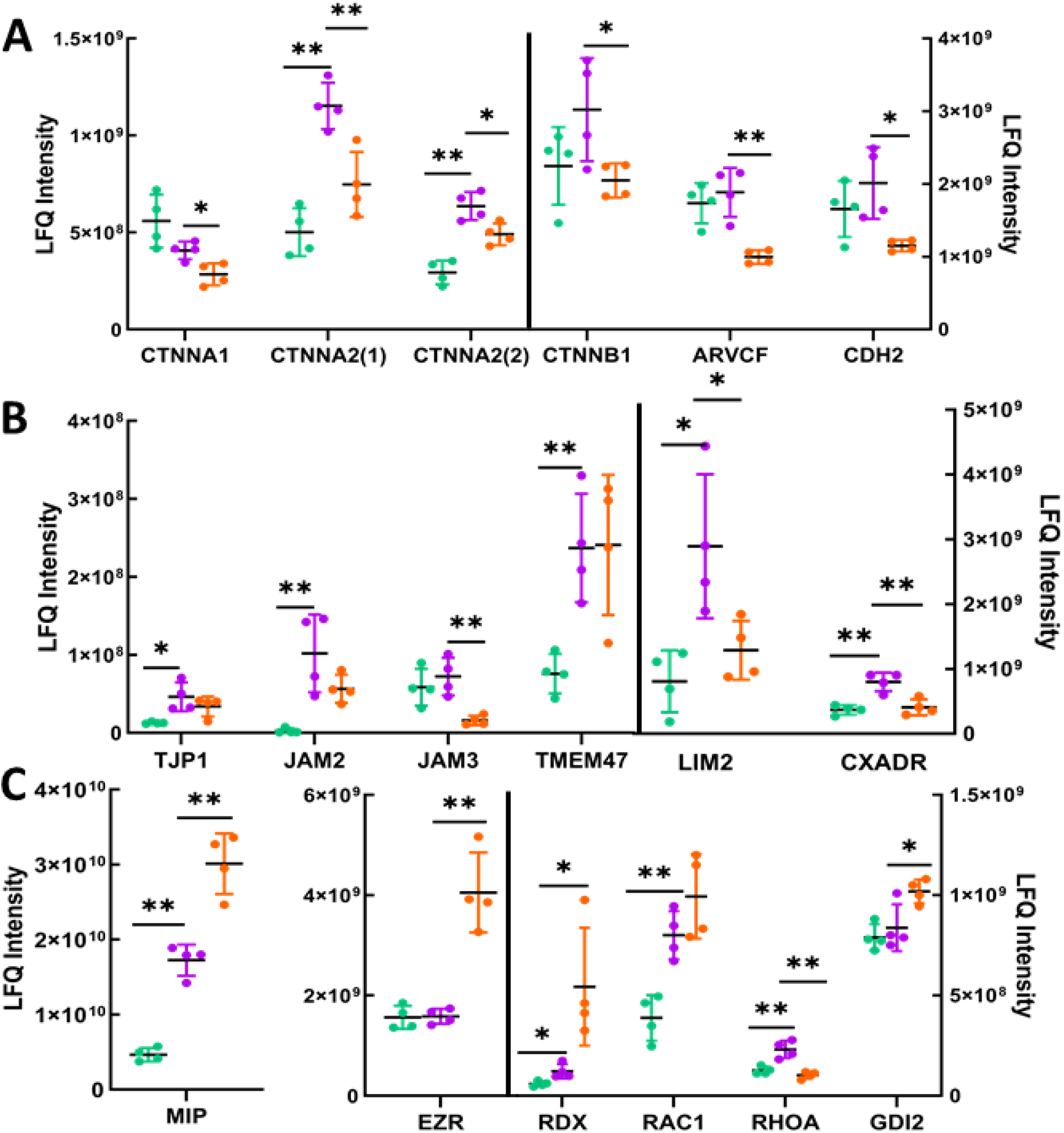
Protein label free quantification (LFQ) intensity across three regions of the lenses. LFQ intensities of major cell junction-related proteins are plotted. Statistical analysis was only done between OC1 and OC2 or OC2 and IC. Green: OC1, Purple: OC2 and orange: IC. * indicates p<0.05 and ** indicates p<0.01.

Another group of proteins that was significantly changed across the three regions is tight junction associated proteins. Several well-known tight junction proteins were detected in lens fiber cells including tight junction protein ZO-1 (TJP1), junctional adhesion molecule 2 (JAM2) and Coxsackievirus and adenovirus receptor homolog (CXADR). These proteins were significantly increased in OC2 samples compared to OC1 samples and tended to decrease in IC samples even though some changes did not reach statistical significance (Figure 4, Figure 7B). Junctional adhesion molecule 3 (JAM3) was also detected and found to be decreased in IC samples. Two PMP-22/EMP/MP20/Claudin Family proteins: lens membrane intrinsic protein (Lim2) and TMEM47 are present in the lens fiber cells (17). Similar to tight junction proteins, Lim2 was found to be significantly increased in OC2 samples and decreased in IC samples. TMEM47 was increased in OC2 samples and maintained similar levels in IC samples. Similarly, gap junction proteins connexin 50 (GJA8) and connexin 46 (GJA3) increased in OC2 samples compared with OC1 samples and did not show significant changes between OC2 and IC regions.

An additional cell junction type, square array junctions, are present in the deep cortex and mature lens fibers and are found to be especially rich in AQP0 and its cleavage products (20, 21, 31). Our results show that AQP0 was significantly increased in OC2 samples compared to OC1 samples (p<0.001) (Figure 4A) and continued increasing in IC samples compared to OC2 samples (p<0.03) (Supplemental Table 1). As shown in Figures 4B and 7C, along with increased AQP0 in ISF samples, ezrin and radixin are also significantly increased in IC samples, both of which have been shown to interact with AQP0 (32). The increased ezrin in the ISF was also accompanied with decreased ezrin in the SF (Supplemental Table 1).

A protein-protein interaction network was constructed using String analysis (27) for proteins that have the same expression trends in each lens region and protein interaction networks that are connected with cell junction proteins were extracted and are shown in Figure 8. This network analysis shows highly connected nodes with junction proteins including proteins that regulate or associate with the actin cytoskeleton and GTPases and their regulators. Adherens junction and tight junction proteins were clustered together through tight junction protein ZO-1 (TJP1) which is also linked to gap junctions through connexins. These junction proteins are connected with the actin cytoskeleton network and the small GTPase signal transduction network (Figures 8A and 8B). Together with the down-regulation of adherens junction and tight junction proteins in the IC region, many actin binding cytoskeleton proteins are down-regulated. The spectrin-ankyrin network is also linked to adherens junction and tight junction network and showed a consistent trend of changing with these junction networks. Figure 8B also shows proteins associated with focal adhesions (cell–matrix adhesions) such as ITGB1, ITGA6, TLN1, FLNC and FLNB decreased in the IC region. The decrease of EPPD complex proteins in the IC region can also be seen in Figure 8B. Among the proteins increased in the IC region, along with AQP0 and ERM proteins, were proteins involved in regulation of the actin cytoskeleton (ARPC1A, ARPC3, MAPK1 and GSN), proteins belonging to the 14-3-3 family (YWHAE, YWHAG and YWHAE), the small GTPase family (RAN, RHEB, RAB6B) and the GTP-binding protein, ARL3.

**Figure 8:**
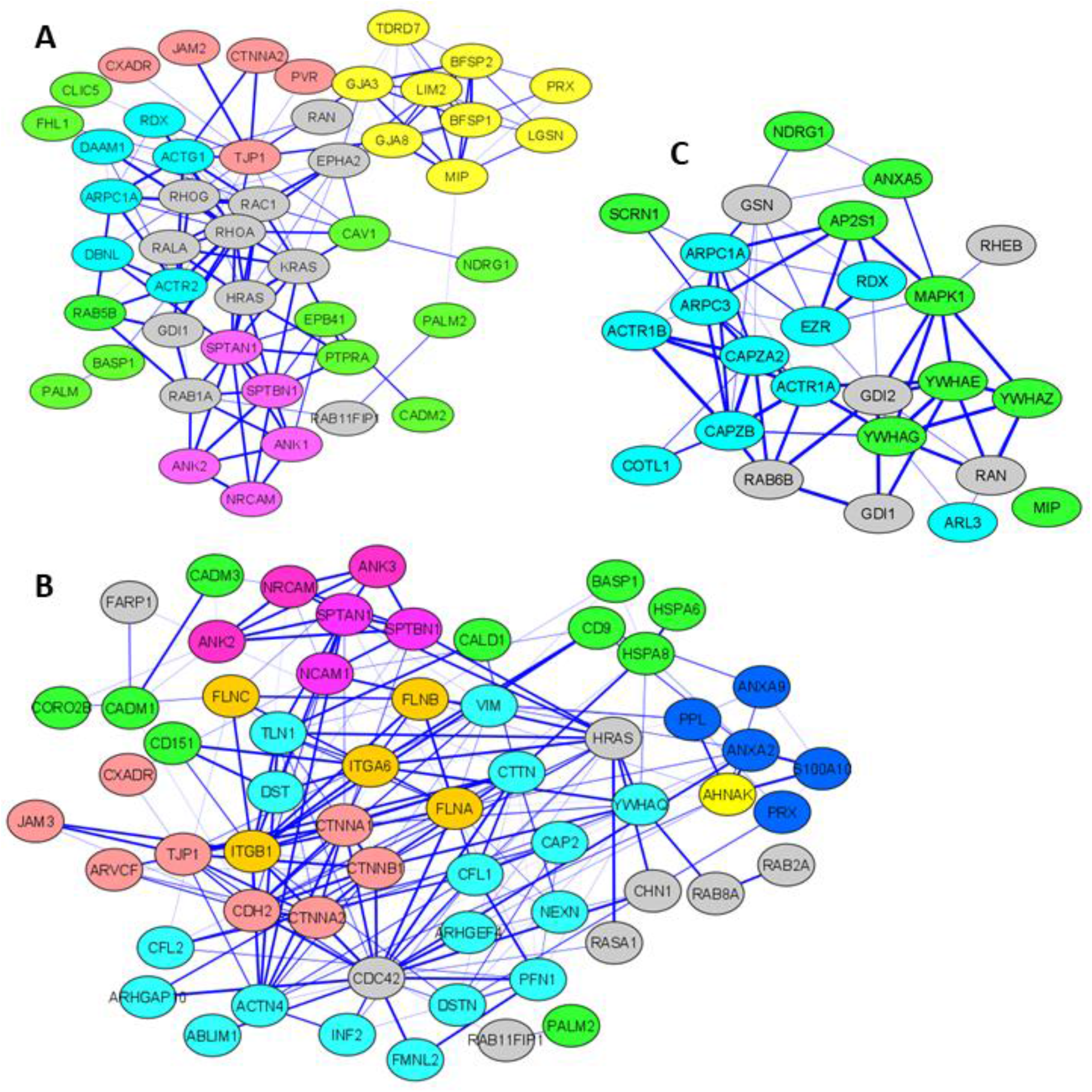
Protein-protein network analysis. Intermolecular interactions of differentially expressed proteins that are associated with cell junction and actin cytoskeleton were generated by STRING 11.0. The minimum required interaction score was medium confidence 0.400. Nodes represent proteins; same node colors indicate membership of the same cluster, and line thickness denotes the confidence level of the interaction. A: proteins increased in OC2 compared with OC1; B: proteins decreased in IC compared with OC2; C: proteins increased in IC compared with OC2.

## Discussion

In this study, we measured differential protein expression across three narrow regions in the cortex of lens equatorial sections to characterize proteins that could be involved in closing lens extracellular space and forming an extracellular diffusion barrier. Our results suggest that the closing of the extracellular space is a two-step process whereby closing of the broad side spacing of fiber cells (OC2 region) is followed by closing of the short side spacing of fiber cells (IC region). Coinciding with changing of extracellular space, cellular morphology and protein expression change dramatically. Cell-cell adhesion proteins are believed to play important roles in regulating the extracellular space and our results demonstrate that junction proteins such as adherens junction proteins, tight junction-related proteins, gap junction proteins and lens fiber membrane intrinsic protein (LIM2) increase in the region where extracellular space between the broad sides of lens fiber cells is reduced. Further experiments are needed to determine whether these proteins are involved in closing the extracellular space of the broad sides of fiber cells.

Based on previous reports, adherens junctions can be found at the narrow faces of the fiber cell especially in the “intersections” where three hexagonal fiber-cells meet (16, 33). Therefore, the increase of adherens junctions in region OC2 is not believed to be involved in the closing extracellular diffusion along the broad sides of the fiber cells. Adherens junctions and associated actin microfilaments are believed to be involved in stabilizing the structural integrity of lens cells during accommodation and in preserving a specific lens shape (29). Gap junctions can be found in both broad sides and narrow sides of the fiber cells (19) and have been reported to restrict the passage of large solutes through extracellular space (9). In addition, connexin 50 can function as an adhesive molecule and Cx50 KO lenses exhibited increased extracellular space (34). Therefore, the increase in gap junction proteins in the OC2 region could contribute the restriction of dye penetration between the broad sides of fiber cells.

Other proteins that could be involved in the narrowing of the broad side extracellular space include LIM2 and several tight junction related proteins including ZO-1, junction adhesion molecule 2 and 3 (JAM 2& JAM3) and coxsackie and adenovirus receptor (CXADR) that are all increased in the OC2 region. The transition of ZO-1 from the narrow faces of the fiber cells in outer cortex to the broad faces of the fiber cells in the inner cortex has been reported (35). Due to the lack of occludins and claudins, tight junctions are not believed to exist in lens fiber cells (14, 16); however, the tight junction protein ZO-1 has been reported to interact with lens connexins (35). In addition, the increase of ZO-1 in the OC2 region coincided with an increase of LIM2, a PMP-22/EMP/MP20/Claudin Family member (17). The cell adhesive property of LIM2 has been reported (36) and insertion of LIM2 into lens fiber cell membranes was found to correlate with the formation of an extracellular diffusion barrier (10). Therefore, LIM2 could contribute to the narrowing of extracellular space; however, whether LIM2 interacts with ZO-1 to form a tight junction like structure is unknown.

In addition to cell-cell adhesion proteins, another important protein that could be involved in closing the extracellular space is lens major membrane protein, AQP0. Our results indicate that AQP0 dramatically increased from the OC1 region to the OC2 region and is one of the very few junction proteins that further increased in the IC region where extracellular diffusion is completely stopped. Previously, AQP0 was found to redistribute throughout the cell membrane in non-nucleated fiber cells from plaque-like structures predominantly on the broad sides in nucleated fiber cells (37). The transition of a certain portion of AQP0 to the narrow sides of the fiber cells could play a key role in closing the extracellular space between the narrow sides since the cell-cell adhesion function AQP0 has been well documented (38, 39).

The localization and distribution of membrane proteins in the plasma membrane are regulated through the membrane-associated cytoskeleton (40). Previously, AQP0 was found to interact with ERM proteins (32), proteins that are linkers of the plasma membrane and the cortical actin cytoskeleton. In this study, ERM proteins ezrin and radixin are among the very few proteins that increase in association with membrane proteins in the IC region. Interactions between AQP0 and ERM proteins could play a role in driving the redistribution of AQP0 in fiber cell membranes. Ezrin is one of the components of the EPPD complex along with periplakin, periaxin and desmoyokin (16). Our results also show periplakin (PPL) and desmoyokin (AHNAK) abundances decreased significantly in the IC region suggesting the decomposition of EPPD complexes. We hypothesize that ezrin released from EPPD complex redistributes in the membrane and interacts with AQP0. A clear transition from soluble fraction to insoluble fraction was detected in the barrier region. Proteins that are involved in this process may also include Rho GTPase and its negative regulators GDP dissociation inhibitors (GDIs). Our results show GDI2 and RAC1 also increased in the IC region. Previously, reduced Rho GTPase activity disrupted the redistribution of AQP0 in lens fiber cell (41, 42). The association of ERM proteins and plasma membrane is regulated by the Rho-dependent signaling pathway (43) and direct interaction of GDI with ERM proteins has been reported (44).

The dramatic increase in expression of lengsin (LGSN), chloride intracellular channel protein 5 (CLIC5), and lactase like protein (LCTL) suggest important roles for these proteins in lens fiber cells. Lengsin was reported to co-localize with lens intermediate filament proteins in maturing fiber cells and may act as a component of the cytoskeleton itself or as a chaperone for the reorganization of intermediate filaments (45). A circulating flux of Cl^-^ ions mediated by an unknown chloride channel is believed to be involved in the operation of the lens microcirculation system and a chloride channel inhibitor causes extracellular dilation (46). We predict the unidentified chloride channel reported by Webb et al. (46) is CLIC5. Even though CLIC1 was also detected, its abundance was significantly decreased in maturing fiber cells. The exchange of chloride channels suggests that CLIC5 must play a particular role in the lens. The function of chloride channels in the lens may include maintenance of a steady hydrated state and cell volume, regulation of pH and maintenance of a circulating flux of Cl^-^ ions (46). LCTL was found to be essential for CLIC5 expression (47). In addition, LCTL is highly expressed in the lens and was reported to regulate suture formation (47). Therefore, despite direct experimental evidence, we hypothesize that these proteins could play important roles in lens microcirculation system.

In addition to barrier related proteome changes, this study also identifies protein changes associated with some important biological processes involved in fiber cell differentiation such as organelle degradation, cytoskeletal organization, and cellular metabolism. These results will be the focus of a separate report.

The current study identifies potential proteins and biological processes that could be involved in regulating the extracellular space of lens fiber cells. Further experiments are needed to confirm the function of these proteins in regulating lens microcirculation system. Furthermore, our current study only focused on a narrow region in the equator of the lens. This study, therefore, does not examine proteome changes between the barrier region and the nucleus region and the proteome changes along the fiber cell length from anterior to posterior poles. Further comparisons between all regions are expected to provide a more complete picture of lens spatially-resolved proteome changes that will help us to understand more about the lens microcirculation system and lens homeostasis.

## Acknowledgements

This work was supported by NIH grants R01 EY013462 and P30 EY008126. Figure captions:

## Supplementary material

Supplemental Figure 1: Pearson correlations between samples

Supplemental Figure 2: Heat map and hierarchical clustering of identified statistically significant changed proteins between different regions in SF

Supplemental Figure 3: Volcano plot of spatially differentially expressed proteins measured in SF.

## Notes

### Competing Interest Statement

The authors have declared no competing interest.

