## Supplemental Figures for "Spatially-Resolved Proteomic Analysis of the Lens Extracellular Diffusion Barrier"

Supplemental Figure 1: Pearson correlations between samples

Pearson correlations of protein signal intensities between samples are shown in A (ISF, OC1 compared with OC2), B (SF, OC1 compared with OC2), C (ISF, OC2 compared with IC) and D (SF, OC2 compared with IC) indicating higher correlation between biological replicates (Green highlighted) and lower correlation between samples from different regions (Yellow highlighted). The correlation coefficients also shows more changes for proteins in insoluble fraction (ISF) from OC2 to IC.

| | | | | | | | | |
|---|---|---|---|---|---|---|---|---|
| OC1_SF_1 | OC1_SF_2 | OC1_SF_3 | OC1_SF_4 | OC2_SF_1 | OC2_SF_2 | OC2_SF_3 | OC2_SF_4 | B |
| | 0.945 | 0.926 | 0.937 | 0.887 | 0.879 | 0.857 | 0.823 | OC1_SF_1 |
| A | | 0.957 | 0.96 | 0.888 | 0.905 | 0.893 | 0.854 | OC1_SF_2 |
| OC1_ISF_1 | | | 0.954 | 0.862 | 0.876 | 0.866 | 0.831 | OC1_SF_3 |
| OC1_ISF_2 | 0.965 | | | 0.873 | 0.889 | 0.871 | 0.843 | OC1_SF_4 |
| OC1_ISF_3 | 0.921 | 0.944 | | | 0.918 | 0.926 | 0.907 | OC2_SF_1 |
| OC1_ISF_4 | 0.903 | 0.913 | 0.911 | | | 0.935 | 0.923 | OC2_SF_2 |
| OC2_ISF_1 | 0.888 | 0.871 | 0.862 | 0.848 | | | 0.931 | OC2_SF_3 |
| OC2_ISF_2 | 0.887 | 0.871 | 0.857 | 0.834 | 0.959 | | | OC2_SF_4 |
| OC2_ISF_3 | 0.867 | 0.866 | 0.868 | 0.851 | 0.949 | 0.933 | | |
| OC2_ISF_4 | 0.821 | 0.814 | 0.816 | 0.825 | 0.911 | 0.894 | 0.92 | |
| | OC1_ISF_1 | OC1_ISF_2 | OC1_ISF_3 | OC1_ISF_4 | OC2_ISF_1 | OC2_ISF_2 | OC2_ISF_3 | OC2_ISF_4 |

| | | | | | | | | |
|---|---|---|---|---|---|---|---|---|
| OC2_SF_1 | OC2_SF_2 | OC2_SF_3 | OC2_SF_4 | IC_SF_1 | IC_SF_2 | IC_SF_3 | IC_SF_4 | D |
| | 0.962 | 0.949 | 0.934 | 0.934 | 0.924 | 0.908 | 0.908 | OC2_SF_1 |
| C | | 0.969 | 0.956 | 0.924 | 0.932 | 0.92 | 0.908 | OC2_SF_2 |
| OC2_ISF_1 | | | 0.954 | 0.92 | 0.935 | 0.926 | 0.913 | OC2_SF_3 |
| OC2_ISF_2 | 0.976 | | | 0.934 | 0.95 | 0.942 | 0.941 | OC2_SF_4 |
| OC2_ISF_3 | 0.959 | 0.942 | | | 0.952 | 0.943 | 0.941 | IC_SF_1 |
| OC2_ISF_4 | 0.91 | 0.883 | 0.91 | | | 0.957 | 0.951 | IC_SF_2 |
| IC_ISF_1 | 0.818 | 0.801 | 0.794 | 0.845 | | | 0.946 | IC_SF_3 |
| IC_ISF_2 | 0.829 | 0.815 | 0.804 | 0.847 | 0.936 | | | IC_SF_4 |
| IC_ISF_3 | 0.808 | 0.775 | 0.811 | 0.859 | 0.91 | 0.895 | | |
| IC_ISF_4 | 0.797 | 0.767 | 0.794 | 0.867 | 0.91 | 0.893 | 0.921 | |
| | OC2_ISF_1 | OC2_ISF_2 | OC2_ISF_3 | OC2_ISF_4 | IC_ISF_1 | IC_ISF_2 | IC_ISF_3 | IC_ISF_4 |

Supplemental Figure 2: Heat map and hierarchical clustering of identified statistically significant changed proteins between different regions in SF

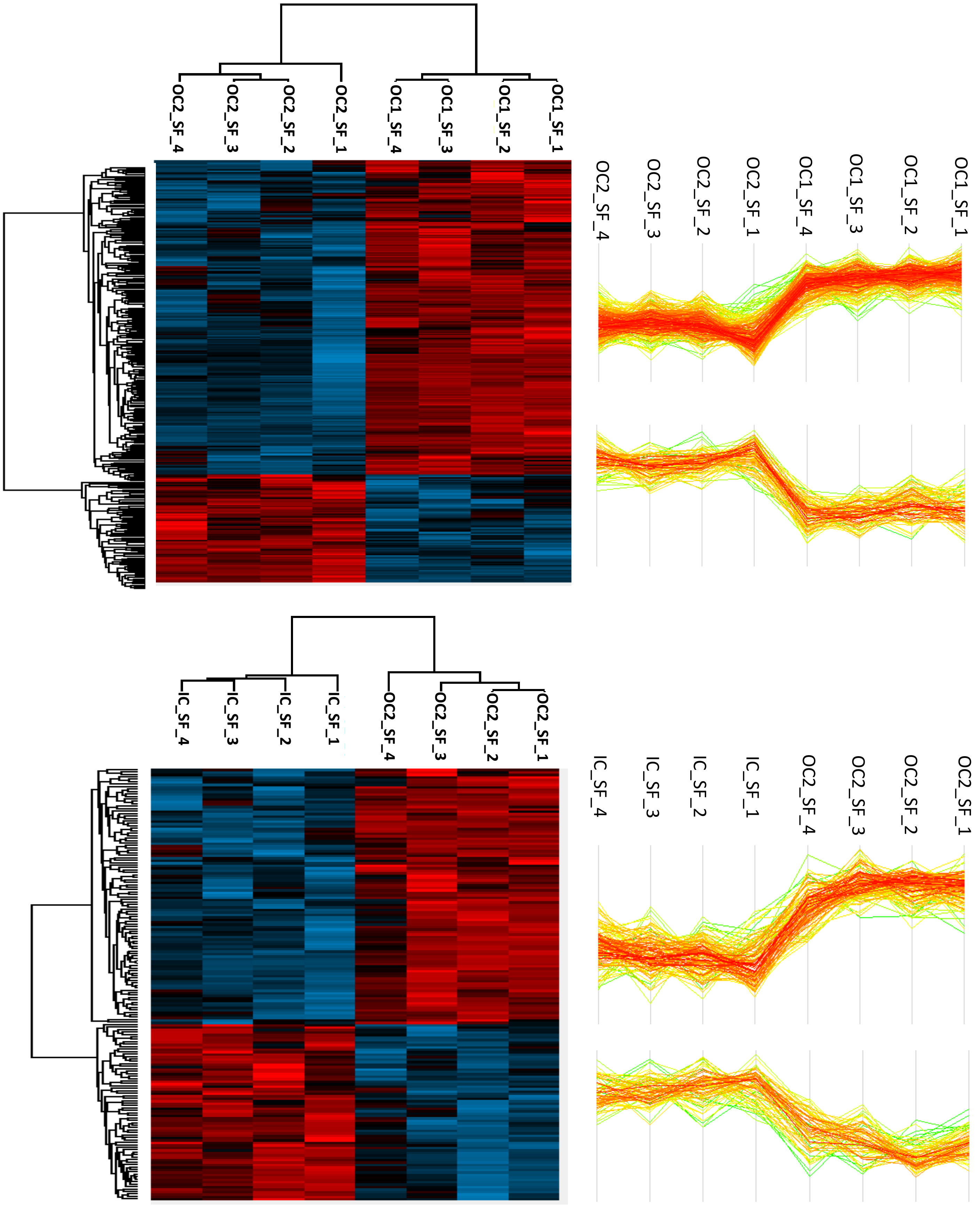

Supplemental Figure 3: Volcano plot of spatially differentially expressed proteins measured in SF. A: comparison between OC2 and OC1; B: comparison between IC and OC2.

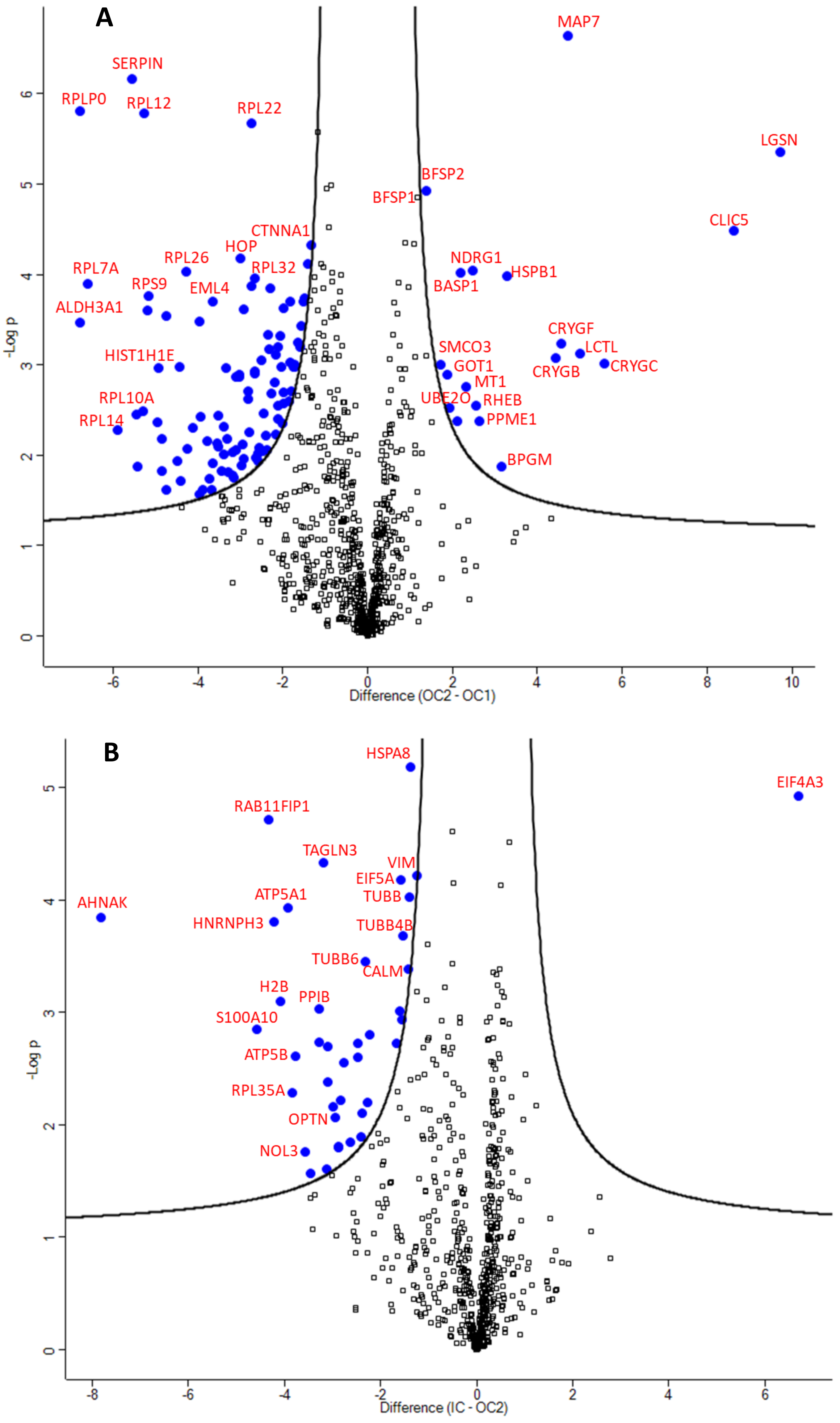
